# All-trans retinoic acid induces synaptopodin-dependent metaplasticity in mouse dentate granule cells

**DOI:** 10.1101/2021.07.05.451170

**Authors:** Maximilian Lenz, Amelie Eichler, Pia Kruse, Julia Muellerleile, Thomas Deller, Peter Jedlicka, Andreas Vlachos

## Abstract

The vitamin A derivative all-trans retinoic acid (atRA) is a key mediator of synaptic plasticity. Depending on the brain region studied, distinct effects of atRA on excitatory and inhibitory neurotransmission have been reported. However, it remains unclear how atRA mediates brain region-specific effects on synaptic transmission and plasticity. Here, we used intraperitoneal injections of atRA (10 mg/kg) in adult male C57BL/6J mice to study the effects of atRA on excitatory and inhibitory neurotransmission in the mouse fascia dentata. In contrast to what has been reported in other brain regions, no major changes in synaptic transmission were observed in the ventral and dorsal hippocampus 6 hours after atRA administration. Likewise, no evidence for changes in the intrinsic properties of hippocampal dentate granule cells was obtained in the atRA-treated group. Moreover, hippocampal transcriptome analysis revealed no essential changes upon atRA treatment. In light of these findings, we tested for the metaplastic effects of atRA, i.e., for its ability to modulate synaptic plasticity expression in the absence of major changes in baseline synaptic transmission. Indeed, *in vivo* long-term potentiation (LTP) experiments demonstrated that systemic atRA treatment improves the ability of dentate granule cells to express LTP. The plasticity-promoting effects of atRA were not observed in synaptopodin-deficient mice, thus extending our previous results on the relevance of synaptopodin in atRA-mediated synaptic strengthening in the mouse prefrontal cortex. Taken together, our data show that atRA mediates synaptopodin-dependent metaplasticity in mouse dentate granule cells.

## INTRODUCTION

Adaptive processes in synaptic sites of the central nervous system are fundamental for normal brain function [1; 2; 3]. Several major signaling pathways and mechanisms that mediate and modulate synaptic plasticity have been identified in the past decades [4; 5; 6]. One of the key mediators of excitatory and inhibitory synaptic plasticity among the plasticity-related signaling molecules is the vitamin A derivative all-trans retinoic acid (atRA) [7; 8; 9; 10; 11].

In work leading to this study, we recently showed that atRA potentiates excitatory postsynapses in human cortical slices prepared from neurosurgical access tissue [12]. This observation is consistent with previous reports suggesting that atRA mediates the accumulation of AMPA receptors at synaptic sites [13; 14]. Moreover, we demonstrated that the presence of the plasticity-related protein synaptopodin, which is an essential component of the calcium ion-storing spine apparatus organelle [15; 16; 17; 18] is required for atRA-mediated strengthening of excitatory neurotransmission in the mouse medial prefrontal cortex. In line with these findings, atRA triggered ultrastructural changes of synaptopodin clusters, spine apparatuses, and dendritic spines in human cortical slices [12].

Thus, both atRA and synaptopodin have been firmly linked to the ability of neurons to express distinct forms of synaptic plasticity [7; 19; 20] and their relevance in homeostatic and associative synaptic plasticity is arguably well established. Recent reports have now implicated atRA and synaptopodin in metaplasticity [21; 22], which refers to the ability of neurons to modify their ability to express synaptic plasticity [4; 23]. Considering the role of the hippocampal formation and more specifically the role of the dentate gyrus in memory acquisition [24] and based on previous work regarding the role of synaptopodin in synaptic plasticity (e.g., [25; 26; 27; 28; 29]), we studied the effects of atRA in synaptic transmission and plasticity and its link to synaptopodin in mouse dentate granule cells.

## MATERIALS AND METHODS

### Ethics statement

All experiments were performed according to the German animal welfare legislation and after positive evaluation by the local authorities (University of Freiburg, AZ G-19/152; Faculty of Medicine at the University of Frankfurt, AZ FU/1131). Animals were kept in a 12 hour light/12 hour dark cycle with access to food and water *ad libitum*. All effort was made to minimize pain or distress of animals.

### Pharmacological treatment

All-trans retinoic acid (atRA; Sigma-Aldrich) was dissolved in DMSO and stored at −20°C until further use. The injection solution was prepared immediately before injection by adding corn-oil to prediluted stocks to achieve a final concentration of 5% DMSO (v/v). Before use, the solution was vortexed briefly. The solution was intraperitoneally injected in adult (C57BL/6J; 6–10 weeks old) male mice at an atRA concentration of 10 mg/kg. Control animals were injected with a vehicle-only solution (5% DMSO in corn oil) but otherwise treated equally. After injection, no overt behavioral changes were observed. Experiments were performed 3–6 hours after intraperitoneal injections.

### Preparation of acute mouse hippocampal slices

Adult mice were anesthetized with ketamine/xylazine (100 mg/kg ketamine and 20 mg/kg xylazine) and rapidly decapitated. Brains were removed and further dissected for the preparations of acute slice of the ventral hippocampus as previously described [30]. For the preparation of acute slices of the dorsal hippocampus, rostral and caudal parts of the brains were removed to ensure a stable coronal sectioning. Brains were immediately transferred to a cooled oxygenated extracellular solution (5°C; 5% CO_2_ / 95% O_2_) containing (in mM): 92 NMDG, 2.5 KCl, 1.25 NaH_2_PO_4_, 30 NaHCO_3_, 20 HEPES, 25 glucose, 2 thiourea, 5 Na-ascorbate, 3 Na-pyruvate, 0.5 CaCl_2_, and 10 MgSO_4_, pH = 7.3 to 7.4 at ~7°C (NMDG-aCSF; [31]). 300 μm tissue sections were cut with a Leica VT1200S vibratome. Slices were transferred to cell strainers with 40 μm pore size placed in NMDG-aCSF at 34°C, and the sodium levels were gradually increased following a protocol as described before [31]. After recovery, slices were maintained for further experimental assessment at room temperature in extracellular solution containing (in mM): 92 NaCl, 2.5 KCl, 1.25 NaH_2_PO_4_, 30 NaHCO_3_, 20 HEPES, 25 glucose, 2 thiourea, 5 Na-ascorbate, 3 Na-pyruvate, 2 CaCl_2_, and 2 MgSO_4_.

### Whole-cell patch-clamp recordings

Dentate granule cells in the suprapyramidal blade of the dentate gyrus were recorded in a bath solution (35°C) containing (in mM): 92 NaCl, 2.5 KCl, 1.25 NaH_2_PO_4_, 30 NaHCO_3_, 20 HEPES, 25 glucose, 2 thiourea, 5 Na-ascorbate, 3 Na-pyruvate, 2 CaCl_2_, and 2 MgSO_4_. Granule cell somata close to the molecular layer of the dentate gyrus were visually identified using an LN-Scope (Luigs & Neumann, Ratingen, Germany) equipped with an infrared dot-contrast and a 40x water immersion objective (Olympus, NA 0.8). Recorded signals were amplified using a Multiclamp 700B amplifier, digitized with a Digidata 1550B digitizer and visualized with the pClamp 11 software package. For recordings of spontaneous excitatory postsynaptic currents (sEPSC) and intrinsic cellular properties, patch pipettes with a tip resistance of 3-5 MΩ were used, contained (in mM): 126 K-Gluconate, 4 KCl, 10 HEPES, 4 MgATP, 0.3 Na2GTP, 10 PO-Creatine, 0.3% (w/v) Biocytin (pH = 7.25 with KOH, 285 mOsm/kg). For sEPSC recordings, dentate granule cells were held at −80 mV in voltage-clamp mode. Intrinsic cellular properties were recorded in current-clamp mode. A pipette capacitance of 2.0 pF was corrected and series resistance was compensated using the automated bridge balance tool of the Multiclamp commander. IV-curves were generated by injecting 1 s square pulse currents starting at −100 pA and increasing in 10 pA steps until +500 pA injection was reached (sweep duration: 2 s). Spontaneous inhibitory postsynaptic currents (sIPSCs) were recorded in the same extracellular solution by adding the AMPA receptor inhibitor CNQX (10 μM, Abcam) and the NMDA receptor inhibitor APV (10 μM, Abcam). Patch pipettes for sIPSC recordings contained (in mM) 40 CsCl, 90 K-gluconate, 1.8 NaCl, 1.7 MgCl_2_, 3.5 KCl, 0.05 EGTA, 2 MgATP, 0.4 Na_2_GTP, 10 PO-Creatine, 10 HEPES (pH = 7.25 with KOH, 290 mOsm) and granule cells were held at −70 mV during the recordings. Series resistance was monitored and recordings were discarded if series resistance reached > 30 MΩ.

### Post-hoc labeling of patched dentate granule cells

Acute slice preparations were fixed in 4% PFA/ 4% sucrose (w/v, phosphate buffered saline, PBS) at room temperature and stored at 4°C overnight in the same solution. After fixation, slices were washed in PBS and consecutively incubated for 1 hour with 10% (v/v) normal goat serum (NGS) in 0.5% (v/v) Triton X-100 containing PBS to reduce unspecific staining. For *post hoc* visualization of patched dentate granule cells, sections were incubated for 3 hours with streptavidin-Alexa Fluor 488 (Invitrogen, #S32354; 1:1000 dilution in 10% (v/v) NGS, 0.1% (v/v) Triton X-100 containing PBS) at room temperature. Sections were washed in PBS and incubated with DAPI for 10 minutes (Thermo Scientific, #62248; 1:5000 dilution in PBS) to visualize cytoarchitecture. After washing, sections were transferred onto glass slides and mounted with fluorescence anti-fading mounting medium (DAKO Fluoromount). Confocal images were acquired using a Leica SP8 laser-scanning microscope equipped with a 20x multi-immersion (NA 0.75; Leica) and a 40x oil-immersion (NA 1.30; Leica) objective. Image stacks were acquired in tile scanning mode with the automated stitching function of the LasX software package.

### RNA isolation and transcriptome analysis

Hippocampi were isolated from the brain of adult mice and immediately transferred into RNA Protection buffer (New England Biolabs) and RNA was consecutively isolated using a column based RNA isolation kit according to the manufacturer’s instructions (Monarch^®^ Total RNA Miniprep Kit; #T2010S New England Biolabs). Strand specific cDNA library preparation from polyA enriched RNA (150 bp mean read length) and RNA sequencing was performed by Eurofins Genomics (Eurofins Genomics Europe Sequencing GmbH, Konstanz, Germany). RNA sequencing was performed using the genome sequencer Illumina HiSeq technology in NovaSeq 6000 S4 PE150 XP sequencing mode. For further analysis .fastq-files were provided. All files contained more than 45 M high quality reads having at least a phred quality 30 (> 90% of total reads).

### In vivo perforant path long-term potentiation

3-month-old male C57BL/6J (Synpo^+/+^) or synaptopodin-deficient animals (Synpo^−/−^; with C57BL/6J genetic background) were kept in a 12 hour light/12 hour dark cycle (Scantainer) with access to food and water *ad libitum*. To achieve stable anesthesia, an initial dose of urethane (1.25 g/kg, in sodium chloride solution) was injected subcutaneously (s.c.); 0.1 g/kg supplemental dose as needed. After stable anesthesia was reached, atRA (10 mg/kg in 5% DMSO) or vehicle-only was intraperitoneally injected (blind to experimenter). The surgery and electrode placement were performed as described previously [32; 33]. Briefly, the mouse was placed in a stereotactic frame (David Kopf Instruments) and local anesthesia with prilocaine (Xylonest 1%, Astra Zeneca, s.c. to the scalp) was applied. Cranial access to the brain was established according to coordinates from the mouse brain atlas (Franklin and Paxinos; stimulation electrode: 2.5 mm lateral to the midline, 3.8 mm posterior to bregma; recording electrode: 1.2 mm lateral to the midline, 1.7 mm posterior to bregma). The ground electrode was placed in the neck musculature. Electrophysiological signals were amplified using a Grass P55 A.C. pre-amplifier (Astro-Med) and digitized at a 10 kHz sampling rate (Digidata 1440A, Molecular Devices). Extracellular stimulation was performed using a STG1004 stimulator (Multichannel Systems). A bipolar stimulation electrode (NE-200, 0.5 mm tip separation, Rhodes Medical Instruments) was lowered 1.5-2.2 mm below the surface of the brain to target the angular bundle of the perforant path. Then a tungsten recording electrode (TM33B01KT, World Precision Instruments) was lowered in 0.1 mm increments while monitoring the waveform of the field excitatory postsynaptic potential (fEPSP) in response to 500 μA test pulses until the granule cell layer was reached (1.7-2.2 mm below the surface). The correct placement of the stimulation electrode in the medial portion of the perforant path was verified electrophysiologically by the latency of the population spike (approximately 4 ms), though the activation of some lateral perforant path fibers could not be excluded. Recordings started a minimum of 3 hours after experimental treatment with atRA or vehicle-only. An input-output curve was generated by 30-800 μA current pulses, repeated 3 times at each intensity, 0.1 ms pulse duration, 60 pulses in total at 0.1 Hz. Recording of perforant path-dentate gyrus (PP/DG)-long-term potentiation (LTP) was performed by applying stimuli with a current intensity set to elicit a 1-2 mV population spike (0.1 Hz, 0.1 ms pulse duration). PP/DG-LTP was induced using a weak theta-burst stimulation (TBS) protocol [34] comprising three series of six trains with six 400 Hz current pulses at double the baseline intensity and pulse duration (with 200 ms interval between trains and 20 s interval between series). Following LTP induction, evoked fEPSPs were recorded for 1 hour using the baseline stimulation parameters.

### Quantification and statistics

RNA-Sequencing data were uploaded to the galaxy web platform (public server: usegalaxy.eu; [35; 36; 37]) and transcriptome analysis was performed using the Galaxy platform in accordance with the reference-based RNA-Seq data analysis tutorial [38]. Briefly, adapter sequences, low quality and short reads were removed via the CUTADAPT tool (Galaxy version 1.16.5). Reads were mapped using RNA STAR (Galaxy version 2.7.6a) with the mm10 Full reference genome (Mus Musculus). For initial assessment of gene expression, unstranded FEATURECOUNT (Galaxy version 2.0.1) analysis was performed from RNA STAR output. Statistical evaluation was performed using DESeq2 (Galaxy version 2.11.40.6+galaxy1) with the treatment as primary factor that might affect gene expression. Genes were considered as differentially expressed if the adjusted p-value was < 0.05. Heat maps were generated based on z-scores of the normalized count table.

Single cell recordings were analyzed off-line using Clampfit 11 of the pClamp11 software package (Molecular Devices). sEPSC and sIPSC properties were analyzed using the automated template search tool for event detection [12]. Input resistance was calculated for the injection of −100 pA current at a time frame of 200 ms with maximum distance to the initial de-/hyperpolarization. Resting membrane potential was calculated as the mean baseline value. AP detection was performed using the input-output curve threshold search event detection tool, and the AP frequency was assessed upon the number of APs detected during the respective current injection time. One cell (control group, ventral hippocampus) was excluded from the analysis of intrinsic membrane properties, since the membrane patch lost its integrity during the recordings. One Synpo^−/−^ animal in the vehicle-only group was excluded from further analysis, since an insufficient response to increasing stimulus intensities could be detected in the input-output curve. Action potential plots in Figures 2 and 4 depict cellular responses until 300 pA current injection. Beyond 300 pA, subsets of granule cells in both groups failed to maintain regular action potential firing. The treatment did not significantly affect action potential frequency beyond 300 pA current injection. *In vivo* perforant path LTP was analyzed using Clampfit 10.2 and custom MATLAB (Mathworks) scripts.

Data were statistically evaluated using GraphPad Prism 7 (GraphPad software, USA). Statistical comparisons were made using the non-parametric Mann-Whitney test. For statistical comparison of XY-plots in whole-cell patch-clamp recordings, we used an RM two-way ANOVA test (repeated measurements/analysis) with Sidak’s multiple comparisons. p-values smaller 0.05 were considered a significant difference. Statistical analysis of fEPSP slope data was performed using the Mann-Whitney test for the three terminal data points. In the text and figures, values represent mean ± standard error of the mean (s.e.m.). U-values were provided for significant results only. * p < 0.05; not significant differences are indicated by ‘ns’.

### Digital Illustrations

Figures were prepared using Photoshop graphics software (Adobe, San Jose, CA, USA). Image brightness and contrast were adjusted.

### Data availability statement

RNA-Sequencing data are accessible from the galaxy web platform via the following link: https://usegalaxy.eu/u/maximilian.lenz/h/transcriptome-analysisatra-6h-vs-controlhippocampus.

## RESULTS

### All-trans retinoic acid (atRA) has no major effects on synaptic transmission and intrinsic cellular properties of dentate granule cells in the dorsal hippocampus

Adult male C57BL/6J mice were injected intraperitoneally with atRA (10 mg/kg) or vehicle-only, and acute coronal slices containing the dorsal hippocampus were prepared 6 hours later. AMPA-receptor-mediated spontaneous excitatory postsynaptic currents (sEPSCs) were recorded from mature granule cells in the suprapyramidal blade of the dentate gyrus (Figure 1A-C). In contrast to neocortical neurons [12], atRA had no apparent effects on the mean sEPSC amplitude, whether half-width or area (Figure 1D). However, a significant increase in sEPSC frequencies was observed in the atRA group (Figure 1E).

**Figure 1:**
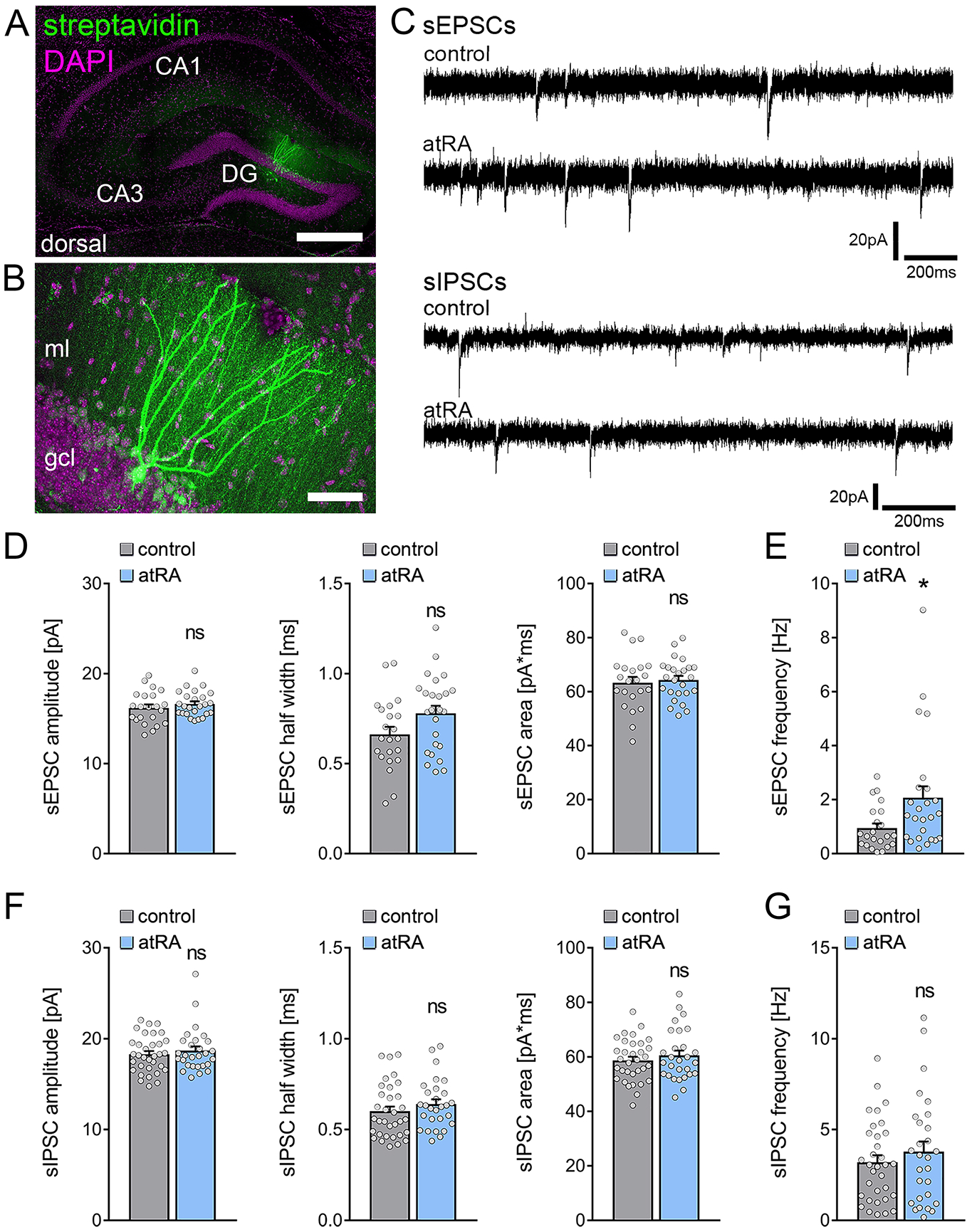
All-trans retinoic acid (atRA) induces no major changes in excitatory and inhibitory neurotransmission onto dentate granule cells of the dorsal hippocampus. (A, B) Example of patched and *post hoc* identified dentate granule cells in acute slices prepared from the dorsal hippocampus. Scale bar (upper panel) = 500 μm; Scale bar (lower panel) = 50 μm. DG, dentate gyrus; gcl, granule cell layer; ml, molecular layer. (C) Sample traces of spontaneous excitatory postsynaptic currents (sEPSCs) and spontaneous inhibitory postsynaptic currents (sIPSCs) recorded from dentate granule cells of atRA (10 mg/kg; i.p.) -treated or vehicle-only (control) animals. (D, E) Group data of sEPSC recordings. A significant increase in sEPSC frequencies is observed. (n_control_ = 22 cells, n_atRA_ = 25 cells in 4 animals; Mann-Whitney test, U_sEPSC frequency_ = 175) (F, G) Group data of sIPSC recordings. (n_control_ = 33 cells, n_atRA_ = 28 cells in 4 animals; Mann-Whitney test) Individual data points are indicated by gray dots. Values represent mean ± s.e.m. (*, p < 0.05; ns, non-significant difference).

Subsequently, we recorded GABA-receptor-mediated spontaneous inhibitory postsynaptic currents (sIPSCs) from dentate granule cells, and we found no significant differences between the two groups (Figure 1C, F, G). The mean sIPSC amplitude—whether half-width or area— and as well as sIPSC frequency were indistinguishable between the two groups. Ostensibly, major changes in inhibitory neurotransmission in the dentate gyrus of atRA-treated mice do not explain the increased sEPSC frequencies (c.f., Figure 1E).

Finally, basic intrinsic properties were assessed (Figure 2). The dentate granule cells from atRA-treated animals were comparable to vehicle-only injected animals. The mean resting membrane potential (Figure 2A) and input resistance (Figure 2B) were similar in both groups. In addition, action potential frequency was not significantly altered in the atRA group in these experiments (Figure 2C). Taken together, these results demonstrate that atRA treatment has no major effects on synaptic neurotransmission and basic intrinsic properties of dentate granule cells. Specifically, atRA does not affect the sEPSC amplitudes of dentate granule cells in the dorsal hippocampus (c.f. [12]).

**Figure 2:**
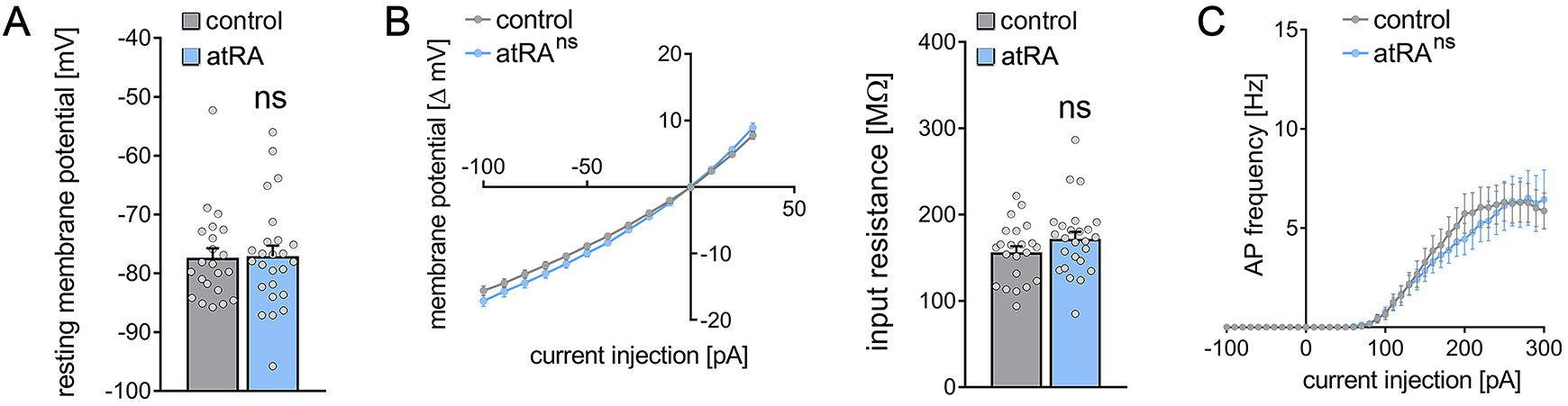
Passive or active membrane properties of dentate granule cells remain unchanged in the dorsal hippocampus following intraperitoneal administration of all-trans retinoic acid (atRA) (A, B) Group data of resting membrane potentials, input-output curves and input resistances. (C) A slight but not significant decrease in action potential (AP) frequencies of dentate granule cells is observed in the atRA group (n_control_ = 22 cells, n_atRA_ = 25 cells in 4 animals each; Mann-Whitney test for column statistics, RM two-way ANOVA followed by Sidak’s multiple comparisons test for action potential frequency analysis)

### All-trans retinoic acid (atRA) has no significant effects on synaptic transmission and intrinsic cellular properties of dentate granule cells in the ventral hippocampus

The ability of neurons to express synaptic plasticity varies along the septotemporal axis of the hippocampus [28; 39; 40; 41]. We therefore tested for the effects of atRA on dentate granule cells in the ventral hippocampus.

A different set of animals was injected intraperitoneally with atRA (10 mg/kg) or vehicle-only, and horizontal slices containing the ventral hippocampus were prepared 6 hours after the injection (Figure 3A). Neither sEPSC nor sIPSC recordings showed any significant differences between the two groups (Figure 3B-D). Likewise, no differences in the active and passive membrane properties were observed (Figure 4). Thus, we concluded that no changes in synaptic transmission and basic intrinsic properties of dentate granule cells are observed in the ventral hippocampus 6 hours after intraperitoneal atRA injection.

**Figure 3:**
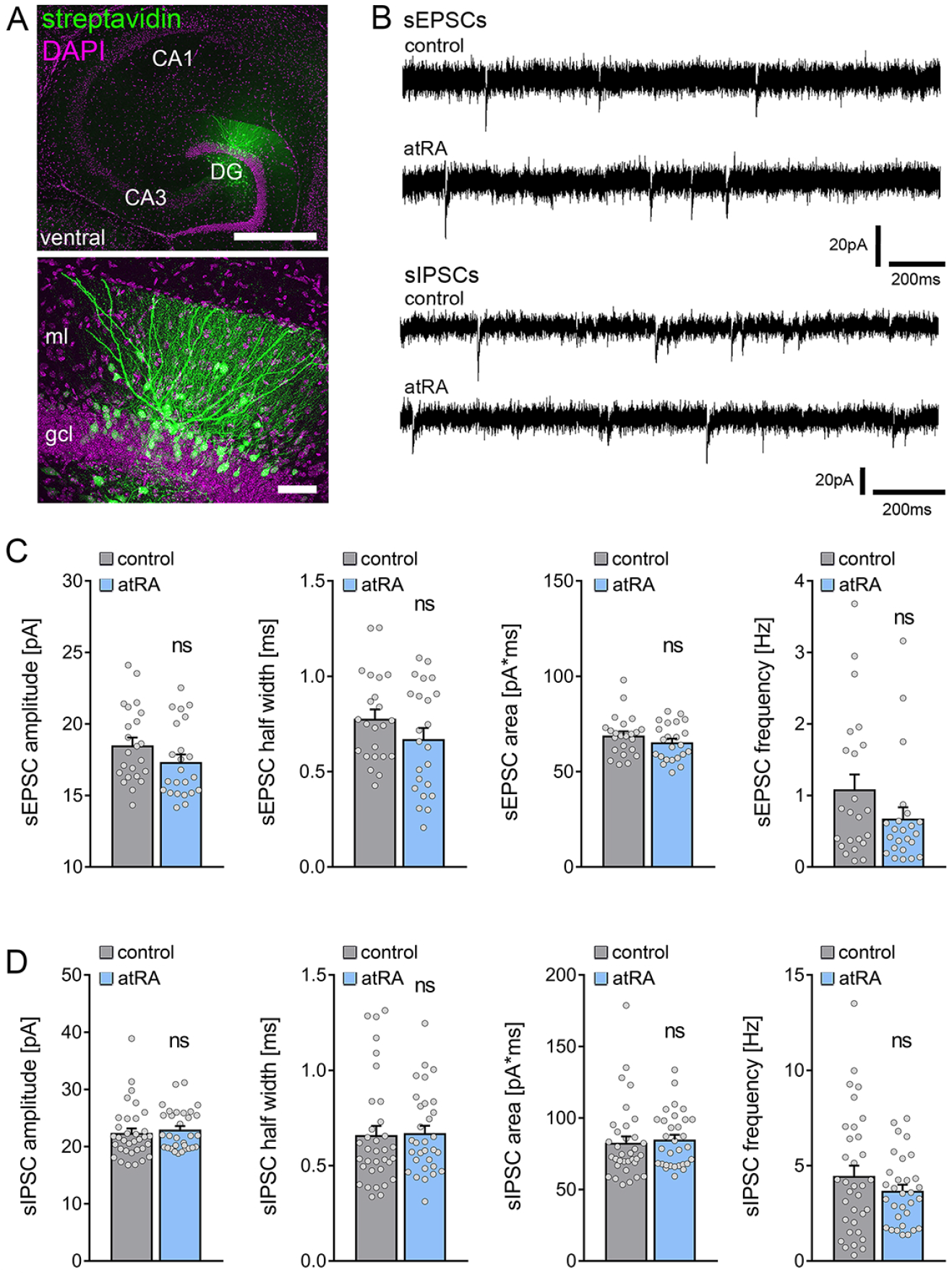
All-trans retinoic acid (atRA) does not induce changes in excitatory and inhibitory neurotransmission of in the ventral hippocampus of adult mice. (A) Example of patched and *post hoc* identified dentate granule cell in acute slices prepared from the ventral hippocampus. Scale bar (upper panel) = 500 μm; Scale bar (lower panel) = 50 μm. DG, dentate gyrus; gcl, granule cell layer; ml, molecular layer. (B) Sample traces of spontaneous excitatory postsynaptic currents (sEPSCs) and spontaneous inhibitory postsynaptic currents (sIPSCs) recorded from dentate granule cells of atRA-treated or vehicle-only (control) animals. (C) Group data of sEPSC recordings (n_control_ = 23 cells, n_atRA_ = 23 cells in 4 animals each; Mann-Whitney test). (D) Group data of sIPSC recordings (n_control_ = 34 cells, n_atRA_ = 31 cells in 4 animals each; Mann-Whitney test). Individual data points are indicated by gray dots. Values represent mean ± s.e.m. (ns, non-significant difference).

**Figure 4:**
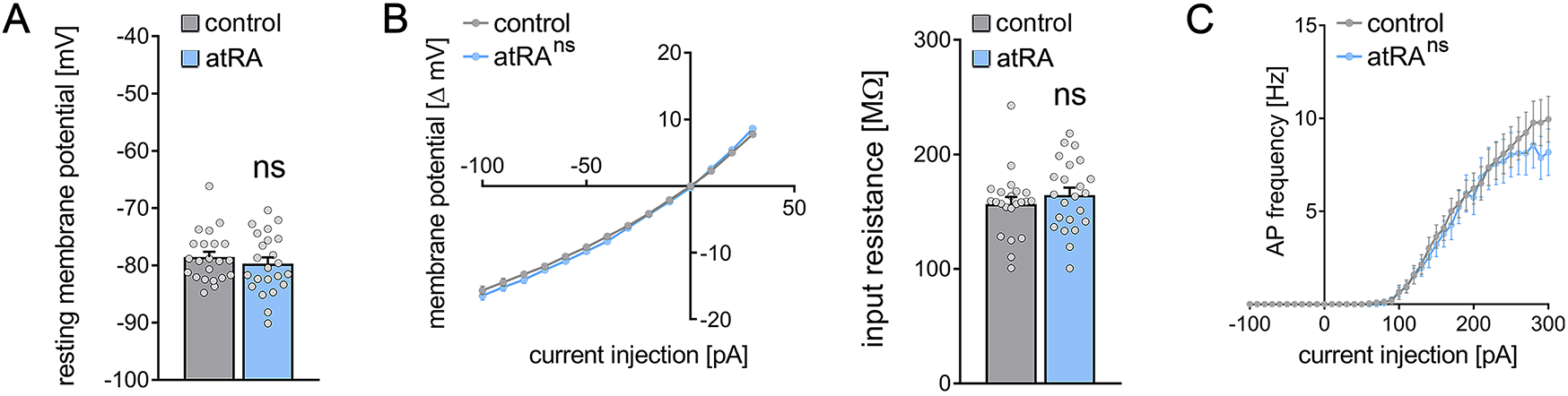
Passive or active membrane properties of dentate granule cells remain unchanged in the ventral hippocampus following intraperitoneal administration of all-trans retinoic acid (atRA) (A-C) Group data of resting membrane potentials (A), input-output curves and input resistances (B) and action potential (AP) frequencies of dentate granule cells in the ventral hippocampus. (n_control_ = 22 cells, n_atRA_ = 23 cells in 4 animals each; Mann-Whitney test for column statistics, RM two-way ANOVA followed by Sidak’s multiple comparisons test for action potential frequency analysis)

### All-trans retinoic acid (atRA) treatment causes only limited changes in the expression of synapse-related genes in the hippocampus

Biological effects of atRA have been reported at the gene transcription level [42]. To further evaluate the effects of atRA in our experimental setting transcriptome analysis was performed in hippocampal tissue samples 6 hours after intraperitoneal atRA or vehicle-only injections (Figure 5). Principal component analysis revealed no major clustering of samples related to the respective treatment (Figure 5A). In line with this observation, only a limited number of significantly regulated genes was identified (29 genes; Figure 5B), representing a z-score heatmap clustering over treatment (Figure 5C). Further analysis of the significantly regulated genes indicated that subsets of these genes relate to atRA-signaling/atRA-metabolism (~ 21%, 6 genes), synaptic transmission (~ 14%, 4 genes), or Wnt signaling (~ 7%, 2 genes), respectively. Of note, the majority of significantly regulated genes did not show any functional clustering (Figure 5D). While these findings demonstrate that intraperitoneally injected atRA reached and affected the hippocampus, no major changes in synaptic genes were detected 6 hours after administration of atRA.

**Figure 5:**
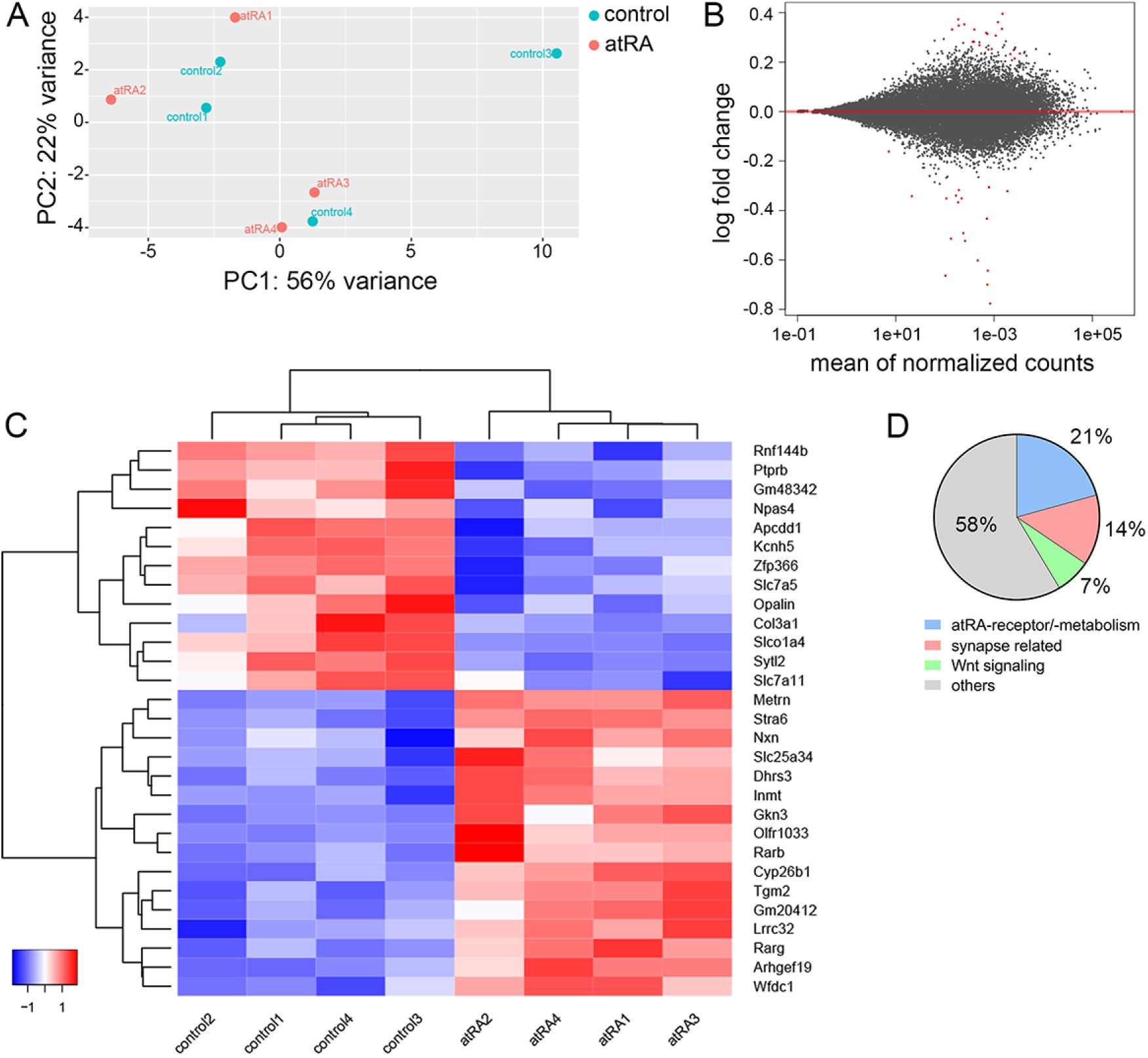
Hippocampal transcriptome analysis reveals no major differences in synapse-related genes following intraperitoneal injection of all-trans retinoic acid (atRA)-treatment. (A) Principal component analysis with “treatment” as primary factor reveals no treatment-specific clustering of hippocampal mRNA samples (n = 4 animals, one hippocampus each). (B) DESeq2-Analysis indicates the differential expression of 29 genes with a moderate |log2FC| < 1 (visualization by MA plot). (C) Heatmap showing the z-scores of differentially expressed genes. The differential expression of genes depend on the atRA-treatment, as indicated by the z-score clustering. (D) Subsets of genes can be attributed to atRA-signaling or atRA-metabolism, synaptic transmission and Wnt-signaling, respectively.

### All-trans retinoic acid (atRA) mediates synaptopodin-dependent metaplasticity in the dentate gyrus

In light of the plasticity-promoting effects of atRA [8; 21; 43], including our recent findings in the mouse and human neocortex [12], we theorized that atRA could induce metaplasticity in the dentate gyrus. Specifically, the lack of essential changes in synaptic transmission and intrinsic cellular properties detected 6 hours after atRA injections prompted the hypothesis, that atRA may modulate the ability of neurons to express synaptic plasticity.

To test for the effects of atRA on the ability of neurons to express synaptic plasticity, long-term potentiation (LTP) experiments on perforant path (PP) synapses were carried out in anesthetized mice (Figure 6A). atRA (10 mg/kg) was injected intraperitoneally 3–6 hours prior to experimental assessment and LTP was probed through electric stimulation of the perforant path with a theta burst stimulation (TBS) protocol (Figure 6A, B). Consistent with our single-cell recordings, which showed no major differences in synaptic and intrinsic cellular properties of dentate granule cells in the atRA group (c.f., Figure 1–4), no significant difference in input-output properties was observed between atRA-treated and vehicle-only animals in these experiments (Figure 6C). Weak TBS was applied to the medial perforant path. As shown in Figure 6D, a significant increase in the fEPSP slopes was observed in both groups upon plasticity induction. However, the increased fEPSP slopes persisted for at least 60 minutes in the atRA-treated mice. We concluded that atRA promotes the ability of dentate granule cells to express long-term synaptic plasticity, while not critically affecting baseline synaptic transmission before plasticity induction.

**Figure 6:**
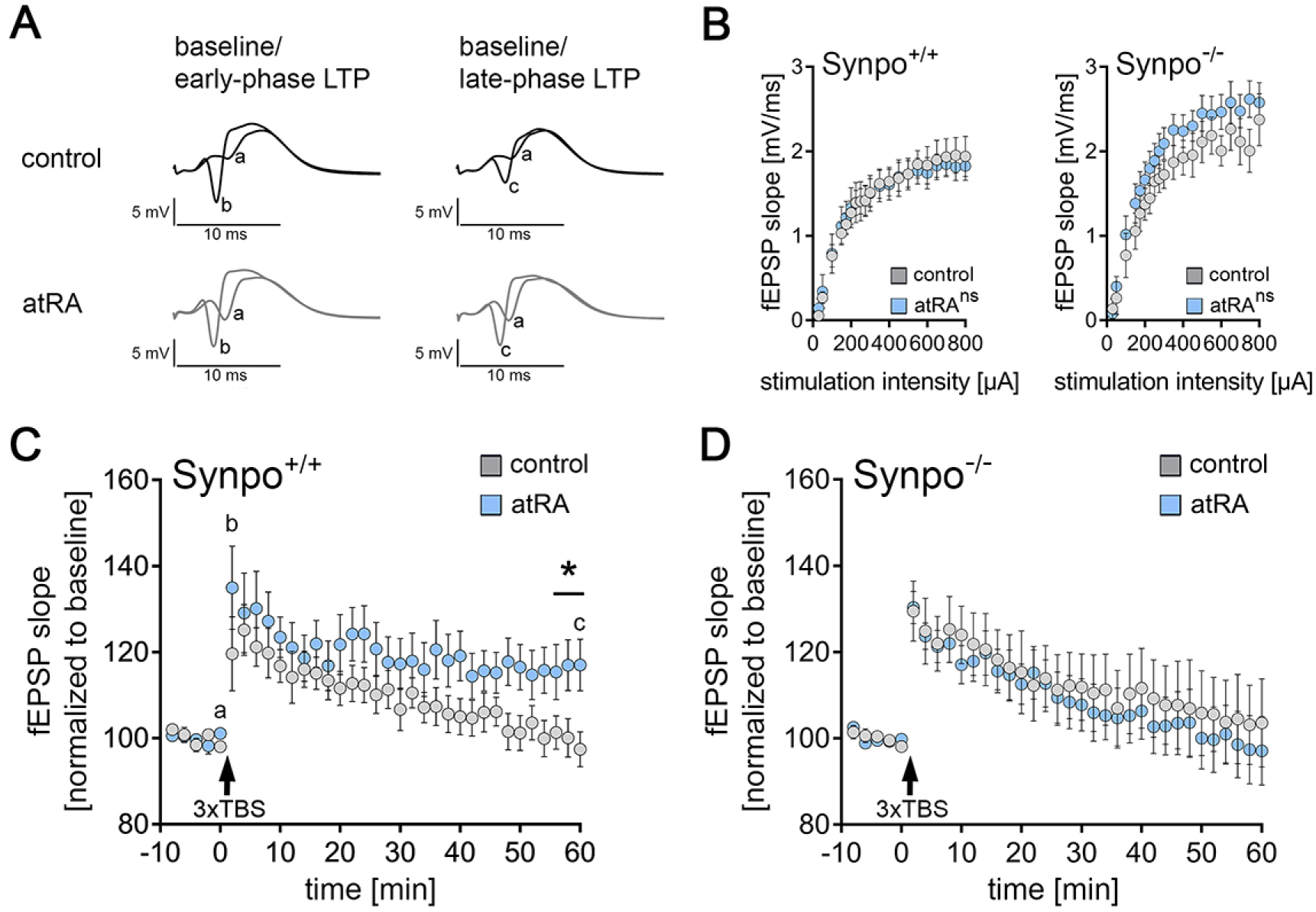
Intraperitoneal injection of all-trans retinoic acid (atRA) improves synaptic plasticity in the dentate gyrus of wildtype but not synaptopodin-deficient mice. (A) *In vivo* long-term potentiation (LTP) experiments on perforant path (PP) synapses were carried out in anesthetized mice using a weak theta-burst stimulation (TBS) protocol. Representative traces of fEPSP recordings in wildtype mice at indicated points in time (a, b, c) after induction of LTP in vehicle-only controls and atRA-injected mice (10 mg/kg, i.p.; 3-6 hours prior to recordings). (B) Input-Output properties of wildtype and synaptopodin-deficient animals (Synpo^+/+^: n_control_ = 9 animals, n_atRA_ = 9 animals; Synpo^−/−^: n_control_ = 7 animals, n_atRA_ = 8 animals. RM two-way ANOVA with Sidak’s multiple comparisons). (C) Group data of fEPSP slopes in wildtype mice (Synpo^+/+^: n_control_ = 9 animals, n_atRA_ = 9 animals; Mann-Whitney test, U = 13-17 for three terminal data points). (D) Group data of fEPSP slopes in synaptopodin-deficient mice (Synpo^−/−^: n_control_ = 7 animals, n_atRA_ = 8 animals; Mann-Whitney test). Values represent mean ± s.e.m. (ns, non-significant difference, * p < 0.05).

To confirm and extend these findings, LTP experiments were carried out in synaptopodin-deficient mice. We have recently demonstrated that synaptopodin is required for atRA-mediated synaptic strengthening in the neocortex [12]. Indeed, atRA- and vehicle-only-treated synaptopodin-deficient animals were indistinguishable in these experiments, and within 60 minutes fEPSP slopes had returned to baseline in both groups (Figure 6C and E). Taken together, we concluded that the intraperitoneal administration of atRA induces metaplastic changes, and that the presence of synaptopodin is required for atRA-mediated metaplasticity.

## DISCUSSION

In this study, we investigated the effects of systemic, i.e., the intraperitoneal application of atRA on synaptic transmission and the plasticity of mature dentate granule cells in the adult hippocampus. Our results demonstrate that atRA promotes the ability of dentate granule cells to express synaptic plasticity. Specifically, a persistent strengthening of excitatory neurotransmission was observed after *in vivo* LTP-induction in the atRA-treated animals. In line with our recent findings, we showed that the presence of synaptopodin is required for atRA-mediated synaptic plasticity [12]. Aside from an increase in sEPSC frequencies in the dorsal hippocampus, atRA had no significant effect on baseline excitatory and inhibitory synaptic transmission in the dentate gyrus. Hence, we propose that atRA modulates the ability of neurons to express synaptic plasticity consistent with a synaptopodin-dependent metaplastic effect of atRA.

Vitamin A metabolites, such as atRA and their related signaling pathways, have been linked to various physiological brain functions, including synaptic plasticity [7; 12; 43; 44]. Accordingly, atRA has been evaluated in disease models and patients with brain disorders associated with cognitive decline, including Alzheimer’s disease, Fragile X syndrome, and depression [11; 45; 46; 47; 48]. However, the precise mechanisms through which atRA asserts its effects on synaptic transmission and plasticity in health and disease are subjects of further investigations.

In a recent study, we demonstrated that atRA induces structural and functional synaptic changes in neurons of the adult human cortex [12]. Specifically, increased sEPSCs amplitudes were observed 6 hours after exposure of acute cortical slices to 1 μM atRA. These functional changes correlated well with increased spine head sizes, and corresponding changes in synaptopodin clusters and spine apparatus organelles were observed [12]. Consistent with these findings, increased sEPSC amplitudes were observed in acute cortical slices prepared from the medial prefrontal cortex of wildtype but not synaptopodin-deficient mice [12]. These findings identified atRA as a potent mediator of synaptic plasticity in the adult human cortex. Furthermore, they suggest that synaptopodin-dependent signaling pathways are involved in mediating the synaptic effects of atRA.

In the present study, however, we did not observe major changes in synaptic neurotransmission onto dentate granule cells in neither the ventral nor the dorsal hippocampus. Specifically, no changes in sEPSC amplitudes were observed [12]. Previous studies reported differential effects of atRA in distinct brain regions. For example, in the visual cortex, a reduction in inhibitory neurotransmission and no effect on excitatory neurotransmission were observed [44], while in the somatosensory cortex, evidence for increased inhibitory synaptic strength in the absence of changes in excitatory neurotransmission was provided [10]. Likewise, in the hippocampal CA1 region, atRA seems to potentiate excitatory while depressing inhibitory synapses [9]. These findings support the notion that atRA may assert its effect on synaptic transmission in a brain region-specific manner. However, differences in the respective experimental settings must be carefully considered, such as the use of distinct tissue preparations (acute slices vs. organotypic tissue cultures vs. dissociated neurons) and differences in atRA administration (systemic vs. local; *in vivo* vs. *ex vivo*), Consequently, more work is required to better understand the distinct effects of atRA on synaptic transmission and to determine for example possible concentration-dependent effects and the impact of a single dose vs. repeated (long-term) administration of atRA.

The results of our mRNA analysis confirmed that intraperitoneally injected atRA affects the hippocampus and leads to transcriptional changes related to atRA-signaling/atRA-metabolism. These findings are in line with previous work demonstrating retinoic acid signaling in the hippocampus [49]. Of note, although a small number of genes showed transcriptional alterations, we did not observe major changes in synaptic or plasticity-related genes. Thus, it appears unlikely that atRA affects plasticity primarily via regulating the transcription of plasticity-related genes. Rather, as has been also suggested by others, atRA-induced changes in mRNA translation could account for the rapid effects of atRA on synaptic plasticity [13; 14]. In line with this interpretation, our previous work revealed that atRA-mediated synaptic changes are not observed when protein synthesis is blocked with anisomycin in mouse and human cortical slices [12], suggesting that atRA modulates protein synthesis and, therefore, the availability of proteins required for the induction of synaptic plasticity. Although these findings do not fully exclude the possibility that atRA-related transcriptional changes influence synaptic plasticity to some extent [50], our data are in line with effects of atRA on local protein synthesis, which plays a major role in different forms of synaptic plasticity, including metaplasticity [51; 52; 53].

It is interesting to theorize that atRA may act as a permissive rather than an instructive plasticity factor in this context. That is to say, atRA may not induce specific changes in excitatory and inhibitory neurotransmission in distinct brain regions but rather act by influencing the ability of neurons to express synaptic plasticity. Thus, the specific outcome of atRA treatment on excitatory and inhibitory neurotransmission may depend on the specific stimuli applied or changes in network activity occurring after atRA administration. Because we did not observe major changes in baseline synaptic transmission in the present study, we were able to test for such permissive, metaplastic effects of atRA. Therefore, *in vivo* LTP was probed with a mild plasticity-inducing stimulus [34]. In the absence of any differences in input-output properties before LTP-induction, atRA promoted the ability of neuron to maintain increased synaptic strength 60 minutes after LTP-induction. These findings are consistent with atRA-mediated metaplasticity.

The plasticity-promoting effects of atRA were not observed in synaptopodin-deficient mice, suggesting that synaptopodin is required for atRA-mediated metaplasticity. These findings are in line with our previous work, which showed that the presence of synaptopodin is required for atRA-mediated synaptic strengthening to occur in the mouse prefrontal cortex [12]. Moreover, we were able to demonstrate that atRA triggers an increase in synaptopodin clusters and spine apparatus sizes in human cortical slices [12]. These findings call for a systematic assessment of atRA-mediated synaptopodin-dependent synaptic plasticity, including assessing specific stimuli, network states and other conditions that may trigger associative and homeostatic changes in excitatory and inhibitory neurotransmission. Whether and how ultrastructural changes of spine apparatus organelles [12; 17; 25] reflect the induction of different forms of synaptic plasticity is currently unknown. Considering that both atRA-signaling and synaptopodin-mediated signaling pathways have been associated to pathological brain states such as Alzheimer’s disease [54; 55; 56; 57; 58] we are confident that a better understanding of atRA-mediated synaptopodin-dependent synaptic plasticity may support the development of novel therapeutic strategies aimed at synaptic plasticity modulation.

## DECLARATIONS

## Acknowledgements

We thank Simone Zenker for technical assistance. The work was supported by Else Kröner-Fresenius-Stiftung (EKFS_#2019_A94 to M.L.) and Deutsche Forschungsgemeinschaft (DFG; CRC1080 to T.D. and A.V.; CRC/TRR167 to A.V.). The Galaxy server that was used for some calculations is in part funded by Collaborative Research Centre 992 Medical Epigenetics (DFG grant SFB 992/1 2012) and German Federal Ministry of Education and Research (BMBF grants 031 A538A/A538C RBC, 031L0101B/031L0101C de.NBI-epi, 031L0106 de.STAIR (de.NBI)).

## Author contributions

Author contributions have been assigned according to CRediT taxonomy. ML: Conceptualization, Validation, Formal Analysis, Investigation, Writing-original draft preparation, Visualization, Project administration, Funding acquisition. AE: Investigation, Formal Analysis. PK: Investigation, Formal Analysis. JM: Investigation, Formal Analysis. TD: Resources, Funding acquisition. PJ: Resources, Investigation, Formal Analysis. Funding acquisition. AV: Conceptualization, Methodology, Resources, Writing-original draft preparation, Supervision, Project administration, Funding acquisition.

